# Single haplotype assembly of the human genome from a hydatidiform mole

**DOI:** 10.1101/006841

**Authors:** Karyn Meltz Steinberg, Valerie A. Schneider, Tina A. Graves-Lindsay, Robert S. Fulton, Richa Agarwala, John Huddleston, Sergey A. Shiryev, Aleksandr Morgulis, Urvashi Surti, Wesley C. Warren, Deanna M. Church, Evan E. Eichler, Richard K. Wilson

## Abstract

An accurate and complete reference human genome sequence assembly is essential for accurately interpreting individual genomes and associating sequence variation with disease phenotypes. While the current reference genome sequence is of very high quality, gaps and misassemblies remain due to biological and technical complexities. Large repetitive sequences and complex allelic diversity are the two main drivers of assembly error. Although increasing the length of sequence reads and library fragments can help overcome these problems, even the longest available reads do not resolve all regions of the human genome. In order to overcome the issue of allelic diversity, we used genomic DNA from an essentially haploid hydatidiform mole, CHM1. We utilized several resources from this DNA including a set of end-sequenced and indexed BAC clones, an optical map, and 100X whole genome shotgun (WGS) sequence coverage using short (Illumina) read pairs. We used the WGS sequence and the GRCh37 reference assembly to create a sequence assembly of the CHM1 genome. We subsequently incorporated 382 finished CHORI-17 BAC clone sequences to generate a second draft assembly, CHM1_1.1 (NCBI AssemblyDB GCA_000306695.2). Analysis of gene and repeat content show this assembly to be of excellent quality and contiguity, and comparisons to ClinVar and the NHGRI GWAS catalog show that the CHM1 genome does not harbor an excess of deleterious alleles. However, comparison to assembly-independent resources, such as BAC clone end sequences and long reads generated by a different sequencing technology (PacBio), indicate misassembled regions. The great majority of these regions is enriched for structural variation and segmental duplication, and can be resolved in the future by sequencing BAC clone tiling paths. This publicly available first generation assembly will be integrated into the Genome Reference Consortium (GRC) curation framework for further improvement, with the ultimate goal being a completely finished gap-free assembly.

## INTRODUCTION

The production of a reference sequence assembly for the human genome was a milestone in biology and clearly has impacted many areas of biomedical research (International Human Genome Sequencing, 2004; McPherson et al., 2001). The availability of this resource allows us to deeply investigate genomic structure and variation at a depth previously unavailable (Genomes Project et al., 2012; Kidd et al., 2008). It is these studies that have helped make clear the shortcomings of our initial assembly models and the difficulty of comprehensive genome analysis. While the current human reference assembly is of extremely high quality and is still the benchmark by which all other human assemblies must be compared, it is far from perfect. Technical and biological complexity lead to both missing sequences as well as misassembled sequence in the current reference, GRCh38 (Church et al., 2011; Eichler et al., 2004; Genovese et al., 2013; International Human Genome Sequencing, 2004; Robledo et al., 2002).

The two most vexing biological problems affecting assembly are 1) complex genomic architecture seen in large regions with highly homologous duplicated sequences and 2) excess allelic diversity (Bailey et al., 2001; Kidd et al., 2008; Korbel et al., 2007; Mills et al., 2006; Zody et al., 2008). Assembling these regions is further complicated due to the fact that regions of segmental duplication are often correlated with copy number variants (CNVs) (Sharp et al., 2005). Regions harboring large CNV segmental duplications have been misrepresented in the reference assembly because assembly algorithms aim to produce a haploid consensus. Highly identical paralogous and structurally polymorphic regions frequently lead to non-allelic sequences being collapsed into a single contig or allelic sequences being improperly represented as duplicates. Because of this complexity, a single, haploid reference is insufficient to fully represent human diversity (Church et al., 2011).

It is critical to build multi-allelic reference assemblies to fully capture human diversity. This begins with the generation of a high-quality single-haplotype primary assembly. The availability of at least one accurate allelic representation at these complex loci facilitates the understanding of the structural diversity (Watson et al., 2013). To enable the assembly of these complex regions, we have developed a suite of resources from CHM1, a DNA source containing a single human haplotype (Fan et al., 2002; Taillon-Miller et al., 1997).

A complete hydatidiform mole (CHM) is an abnormal product of conception in which there is a very early fetal demise and overgrowth of the placental tissue. The majority of CHMs are androgenetic and contain only the paternally and X retained derived chromosomes. The phenotype is thought to be a result of abnormal parental contribution leading to aberrant genomic imprinting (Hoffner and Surti, 2012). The absence of allelic variation of the CHM makes it an ideal candidate for producing a single haplotype representation of the human genome. There are a number of existing resources associated with the “CHM1” sample, including a BAC library with end sequences (https://bacpac.chori.org/), an optical map and a BioNano map, some of which have previously been used to improve regions of the reference human genome assembly.

A BAC library constructed from CHM1 DNA (CHORI-17) has been utilized to resolve several very difficult genomic regions, including human-specific duplications at the *SRGAP2* loci on chromosome 1 (Dennis et al., 2012). Additionally, the CHM1 BAC clones were used to generate single haplotype assemblies of regions that were previously misrepresented because of haplotype mixing (Watson et al., 2013) or use of clonal material derived from white blood cells. Both of these efforts contributed to the improvement of the GRCh38 reference human genome assembly, adding hundreds of kilobases of sequence missing in GRCh37, in addition to providing an accurate single haplotype representation of complex genome regions.

Because of the previously established utility of sequence data derived from the CHM1 resource, we wished to develop a complete assembly of a single human haplotype. To this end, we produced a short read-based (Illumina) reference-guided assembly of CHM1 with integrated high quality finished BAC sequences to further improve the assembly. This assembly has been annotated using the NCBI annotation process and has been aligned to other human assemblies in GenBank, including both GRCh37 and GRCh38. Here we present evidence that the CHM1 genome assembly is a high quality draft with respect to gene and repetitive element content as well as a comparison to other reference assemblies. We will also discuss current plans for developing a fully finished genome assembly based on this resource.

## RESULTS

We generated an assembly of the complete hydatidiform mole, CHM1, genome comprised of 23 chromosomes (1-22 and X and MT) with a total sequence length of 3.04 Gb. Contig N50 length is approximately 144 Kbp and scaffold N50 length is 50 Mbp (Table 1). These N50 statistics were based upon the reference guided assembly with BAC tiling paths incorporated. Compared to other WGS human assemblies, HuRef (J. Craig Venter assembly; Genbank GCA_000002125.2), ALLPATHS (Genbank AEKP00000000.1) and YH_2.0 (Genbank GCA_000004845.2), the CHM1_1.1 assembly has a lower contig number and a higher contig N50 demonstrating that CHM1_1.1 is more contiguous than previously generated individual genome assemblies (Figure 1). We incorporated high quality sequence from 382 BAC clones to improve the assembly in complex regions where the GRCh37 reference was incorrect (Figure 2; Table SI).

**Table 1.** Assembly statistics

|  |  |
| --- | --- |
| Total sequence length | 3,037,866,619 |
| Total assembly gap length | 210,229,812 |
| Gaps between scaffolds | 225 |
| Number of scaffolds | 163 |
| Scaffold N50 | 50,362,920 |
| Number of contigs | 40,828 |
| Contig N50 | 143,936 |
| Total number of chromosomes | 23 |

**Figure 1.**
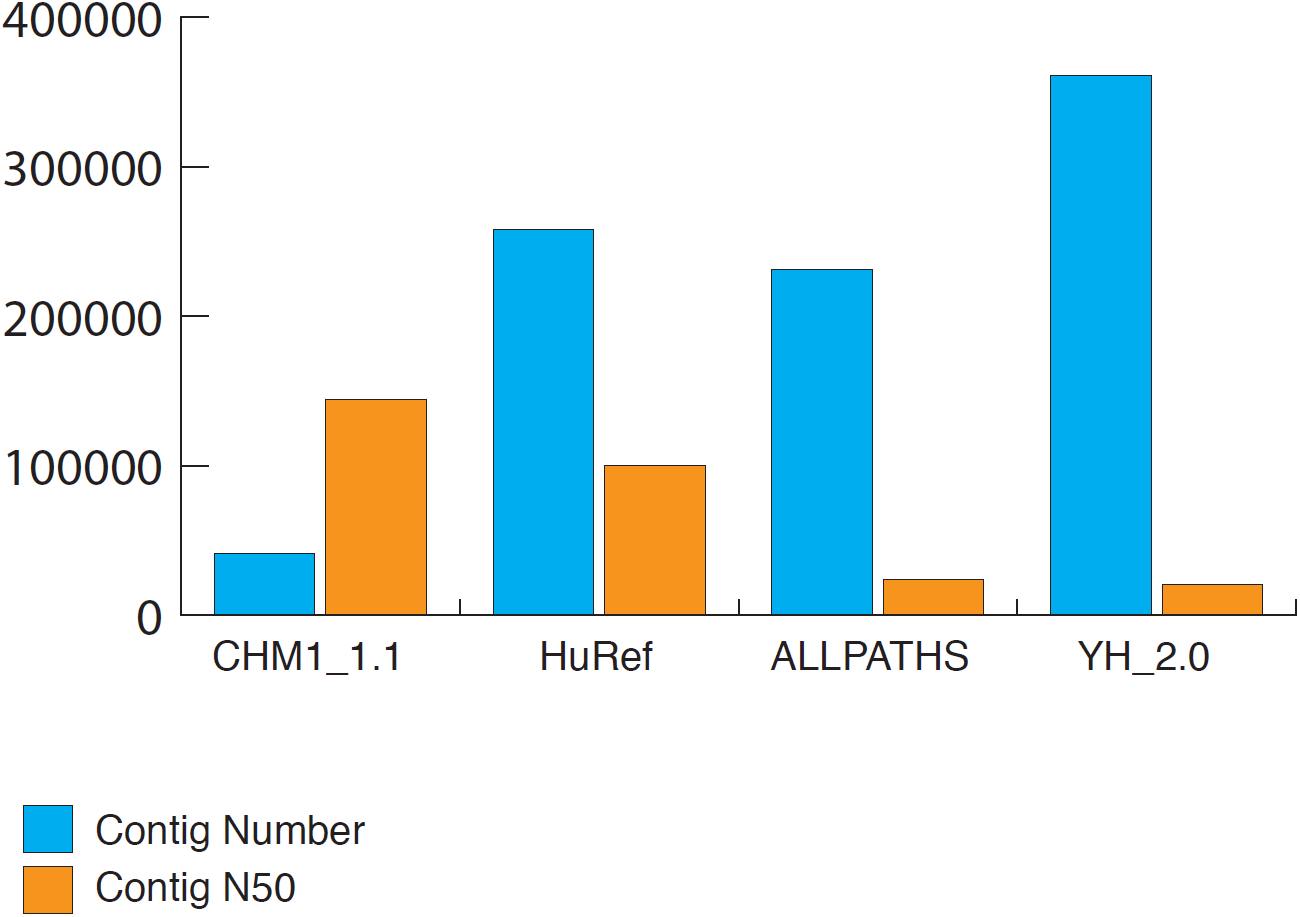
Comparison of contig count and contig N50 between CHM1_1.1 and HuRef, ALLPATHS and YH_2.0 WGS assemblies. CHM1_1.1 has only 10-20% the number of total contigs as the other assemblies and has a contig N50 1.5-6 times larger.

**Figure 2.**
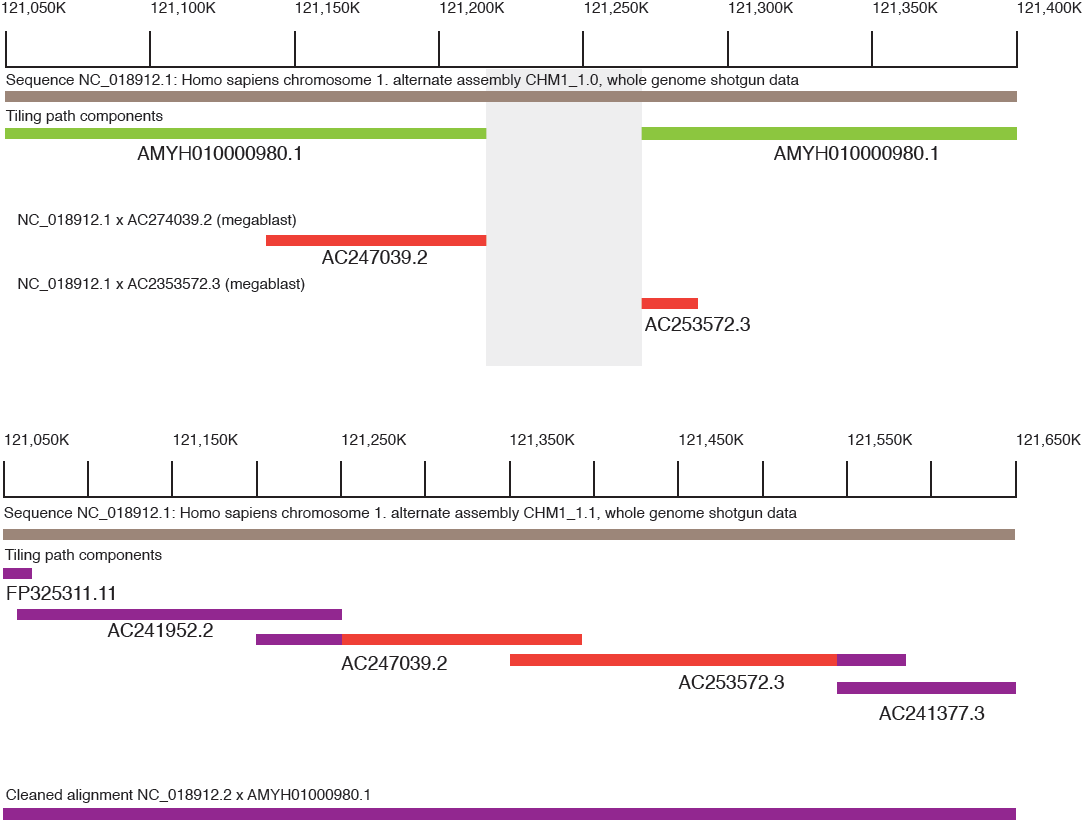
WGS assembly from the first pass (CHM1_1.0; GCF_000306695.1) on chromosome lpl2 (NC_018912.1:121,050,000-121,400,000) demonstrated a gap in the assembly. Using megablast, two CH17 clones (AC247039.2 and AC253572.3) aligned to the region and appeared to span the gap. By incorporating these BAC sequences into the assembly, the gap was subsequently resolved in CHM1_1.1 (NC_018912.2: 121,050,000-121,650,000).

We assessed the integrity and fidelity of CHM1 with respect to the reference by analyzing CHORI-17 BAC end sequencing mapping to GRCh37. Approximately 95.5% percent of clone ends mapped uniquely concordantly, 4% mapped uniquely discordantly and the remaining 0.5% mapped to multiple locations. These statistics indicate that the genomic DNA derived from the CHM1 cell line that was used to create the BAC library and Illumina libraries is not grossly rearranged and represents a suitable template for a platinum reference. In addition, analyses from an optical map generated using CHM1 genomic DNA do not show an excess of structural variants that would suggest somatic rearrangement (Teague et al., 2010). SNP genotyping also confirms the haploid content of the cell line, and karyotyping was performed at several stages during passaging to ensure the integrity (Fan et al., 2002).

## Assessment of assembly quality

### Repeat Content

The assembly was masked with both WindowMasker (Morgulis et al., 2006) and RepeatMasker (Smit et al., 1996-2010) and 34.29% and 47.21% of the assembly was masked, respectively. This is comparable to the repetitive content of GRCh37 (34.24% and 47.15%, WindowMasker and RepeatMasker, respectively). When the repetitive elements are parsed out by type, the numbers of each element are comparable between GRCh37 and CHM1_1.1 (Table 3).

**Table 2.** List of pathogenic/risk alleles in CHM1_1.1. ND = not determined and ND* = not determined, stop gain

| dbSNP137<br>rsID | GRCh37<br>accession | GRCh37<br>chr | GRCh37 pos | CHM1_1.1<br>accession | CHM1_1.1 pos | Risk<br>Allele | Phenotype | Global MAF |
| --- | --- | --- | --- | --- | --- | --- | --- | --- |
| rs1801265 | NC_000001.10 | chr1 | 98,348,885 | NC_018912.2 | 98,464,491 | A | Dihydropyrimidine dehydrogenase deficiency | 0.23 |
| rs12021720 | NC_000001.10 | chr1 | 100,672,060 | NC_018912.2 | 100,788,351 | C | Intermediate maple syrup urine disease type 2 | 0.1 |
| rs121434375 | NC_000001.10 | chr1 | 145,415,341 | NC_018912.2 | 147,387,798 | T | Hemochromatosis type 2A | ND* |
| rs74315325 | NC_000001.10 | chr1 | 145,416,320 | NC_018912.2 | 147,386,819 | A | Hemochromatosis type 2A | ND |
| rs11558492 | NC_000001.10 | chr1 | 231,408,091 | NC_018912.2 | 232,681,743 | G | Rhizomelic chondrodysplasia punctata type 2 | 0.14 |
| rs61744404 | NC_000002.11 | chr2 | 233,390,199 | NC_018913.2 | 233,396,288 | G | Microphthalmia isolated 6 | 0.02 |
| rs1494558 | NC_000005.9 | chr5 | 35,861,068 | NC_018916.2 | 35,863,764 | C | Severe combined immunodeficiency autosomal recessive T cell-negative B cell-positive NK cell-positive | 0.63 |
| rs1494555 | NC_000005.9 | chr5 | 35,871,190 | NC_018916.2 | 35,873,884 | A | Severe combined immunodeficiency autosomal recessive T cell-negative B cell-positive NK cell-positive | 0.66 |
| rs121909192 | NC_000005.9 | chr5 | 69,372,372 | NC_018916.2 | 69,337,143 | C | Spinal muscular atrophy modifier of | ND |
| rs820878 | NC_000005.9 | chr5 | 73,981,270 | NC_018916.2 | 73,413,971 | C | Sandhoff disease infantile type | 0.02 |
| rs119456965 | NC_000005.9 | chr5 | 138,386,649 | NC_018916.2 | 137,819,189 | A | Marinesco-Sjogren syndrome | ND* |
| rs11739136 | NC_000005.9 | chr5 | 169,810,796 | NC_018916.2 | 169,243,672 | T | Hypertension diastolic resistance to | 0.1 |
| rs1800451 | NC_000010.10 | chr10 | 54,531,226 | NC_018921.2 | 54,813,023 | T | Mannose-binding protein deficiency | 0.05 |
| rs10509305 | NC_000010.10 | chr10 | 70,645,376 | NC_018921.2 | 70,927,001 | C | Preeclampsia/eclampsia 4 | 0.14 |
| rs1801252 | NC_000010.10 | chr10 | 115,804,036 | NC_018921.2 | 116,087,941 | G | Resting heart rate | 0.17 |
| rs1169305 | NC_000012.11 | chr12 | 121,437,382 | NC_018923.2 | 121,406,209 | G | Maturity-onset diabetes of the young type 3 | 0.01 |
| rs1154510 | NC_000012.11 | chr12 | 122,295,335 | NC_018923.2 | 122,262,688 | C | 4-Alpha-hydroxyphenylpyruvate<br>hydroxylase deficiency | 0.13 |
| rs2238472 | NC_000016.9 | chr16 | 16,251,599 | NC_018927.2 | 16,335,897 | T | Pseudoxanthoma elasticum | 0.2 |
| rs4784677 | NC_000016.9 | chr16 | 56,548,501 | NC_018927.2 | 57,955,514 | T | Bardet-Biedl syndrome 2 | 0.01 |
| rs4792311 | NC_000017.10 | chr17 | 12,915,009 | NC_018928.2 | 12,923,749 | A | Prostate cancer hereditary 2 | 0.2 |
| rs5911 | NC_000017.10 | chr17 | 42,453,065 | NC_018928.2 | 42,688,697 | C | Bak platelet-specific antigen | 0.4 |
| rs6504649 | NC_000017.10 | chr17 | 48,437,456 | NC_018928.2 | 48,501,299 | G | Pseudoxanthoma elasticum modifier<br>of severity of | 0.27 |

**Table 3.** Comparison of repetitive elements in GRCh37 and CHM1_1.1

|  | **CHM1** | **GRCh37** |
| --- | --- | --- |
| DNA | 482,783 | 461,751 |
| LINE | 1,511,690 | 1,498,690 |
| LTR | 716,720 | 717,656 |
| Low complexity | 376,835 | 371,543 |
| RNA | 778 | 729 |
| SINE | 1,757,213 | 1,793,723 |
| Satellite | 3,099 | 9,566 |
| Simple repeat | 398,210 | 417,913 |
| rRNA | 1,749 | 1,769 |
| scRNA | 1,301 | 1,340 |
| snRNA | 4,306 | 4,386 |
| srpRNA | 1,665 | 1,481 |
| tRNA | 1,852 | 2,002 |
| Other | 19,338 | 15,581 |

**Figure 3.**
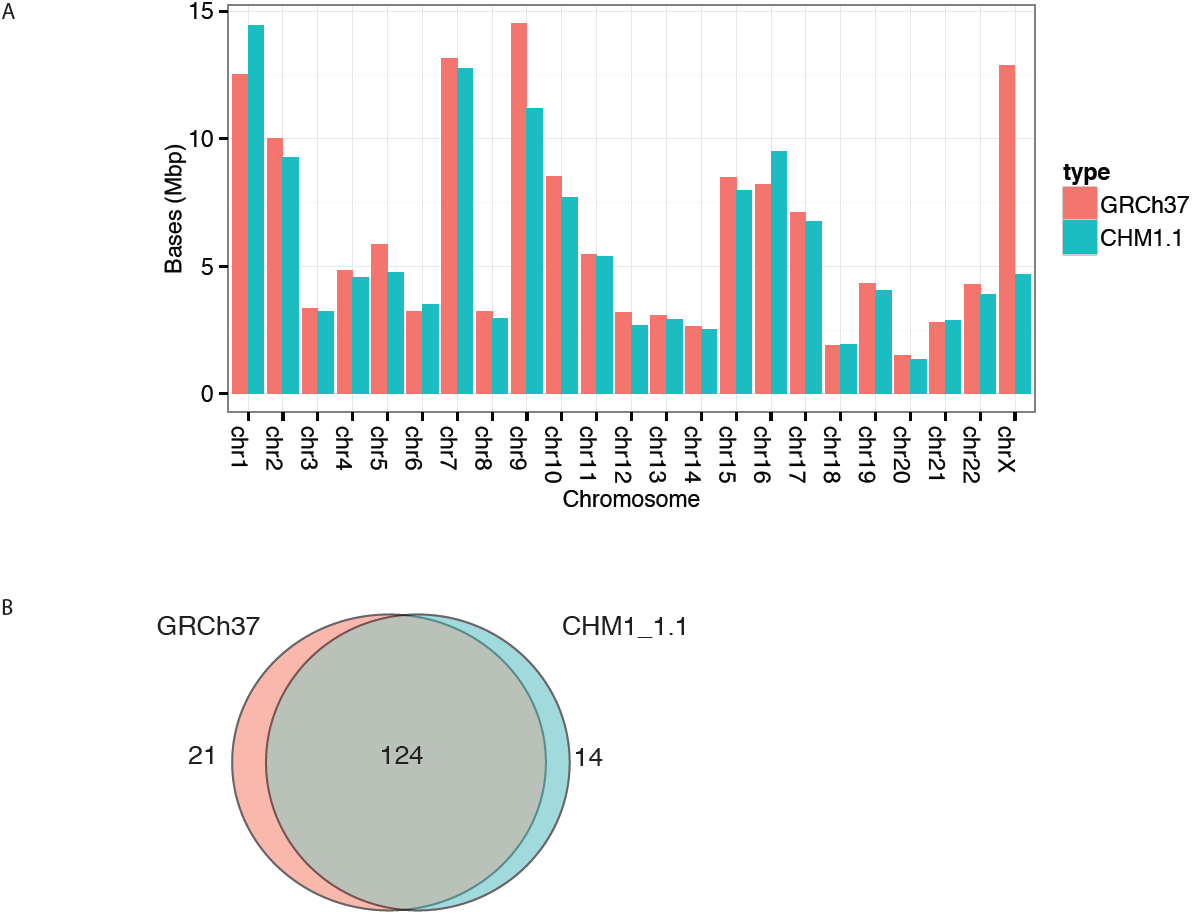
Comparison of segmental duplications in GRCh37 and CHM1.1 assemblies predicted by WGAC analysis by chromosome. Venn-diagram of segmental duplications in GRCh37 and CHM1_1.1 assemblies shows that most duplications are shared between the assemblies..

### Gene Content

This analysis is based on NCBI *Homo sapiens* annotation run 105 (http://www.ncbi.nlm.nih.gov/genome/annotation_euk/Homo_sapiens/105/) that includes GRCh37.p13, CHM1_1.1, HuRef and the single chromosome assembly CRA_TCAGchr7v2. For our comparison, we only used the annotations on the original GRCh37 Primary Assembly sequences, as many of the fix patches in patch release 13 are based on CHM1. Using this annotation run provides a better comparison than the original GRCh37 annotation as the same software and evidence set was used.

Using gene annotation as a proxy for assembly quality, the results indicate that the CHM1_1.1 assembly (39,009 total genes, 19,892 protein coding genes; Table S2) is of higher quality than the HuRef assembly (38,070 total genes, 19,668 protein coding genes), though not quite as good as the GRCh37 assembly (39,947 total genes, 20,072 protein coding genes). The alignment evidence used to support each gene model supports this conclusion. CHM1_1.1 has 21 genes annotated with a ‘transcription discrepancy’ compared to 15 in GRCh37. Interestingly, some genes are problematic in both assemblies, such as MUC8 and MUC19, suggesting that even in a single haplotype background, complex gene family regions can be difficult to assemble. (Supplemental Data: Gene Annotation).

While GRCh37 may have better global gene annotation metrics, there are regions in which CHM1_1.1 performs better. For example, we identified 549 genes unique to the CHM1 assembly (i.e. absent from the GRCh37.p13 primary assembly; Table S3). *MUC3B,* a membrane bound mucin that maps to chromosome 7q22 (|NC_018918.2:100477710-100541651) is annotated only on CHM1_1.1 as predicted from Gnomon gene models. The protein produced by *MUC3B* functions as a major glycoprotein component of mucus gel at the intestinal surface that provide a barrier against foreign particles and microbial organisms. It is part of a tandem duplication involving *MUC3A,* and *MUC3B* is expressed exclusively in the small intestine and colon (Kyo et al., 2001; Pratt et al., 2000). Variants of *MUC3A* have been associated with Inflammatory Bowel Disease, and upregulation of *MUC3* inhibited adherence of pathogenic *E. coli* in human intestinal cells (Pan et al., 2013). The CHM1 version of *MUC3B* contains 4 copies of the tandem repeat.

Other clinically relevant CHM1 genes not present in the GRCh37.p13 primary assembly include *KCNJ18* and *DUX4L. KCNJ18* is a member of a large gene family of potassium inwardly rectifying channels located on 17qll.2 (NC_018928.2: 21605469-21617558). It is expressed mostly in skeletal muscle and regulated by thyroid hormone. Mutations in this gene have been associated with thyrotoxic hypokalemic periodic paralysis (MIM 613239) (Ryan et al., 2010). *DUX4L* encodes a transcription factor comprised of two homeobox domains located within a macrosatellite repeat in the subtelomeric region of 4q (NC_018915.2:190981943-190983264) (Hewitt et al., 1994). Repeat copy number variation is associated with facioscapulohumeral muscular dystrophy (MIM 158900) (Bosnakovski et al., 2008). Both of these genes are now annotated in GRCh38 with information from the CHM1 data.

### Clinical allele analysis

Using data from the NHGRI GWAS catalog and ClinVar, we assessed the number of risk alleles present in the CHM1 genome. Most loci could be successfully remapped from GRCh37 to CHM1 (7962/7991 NHGRI GWAS loci and 43,614/48,516 ClinVar loci) using the NCBI Remap tool (http://www.ncbi.nlm.nih.gov/genome/tools/remap). The CHM1 genotype matched the "risk" allele at 3,284 loci from NHGRI GWAS and 291 loci from ClinVar. CHM1 carries an associated allele for 366 unique phenotypes out of a total of 1089 unique phenotypes in NHGRI GWAS. Of the 291 matching ClinVar alleles, 22 are categorized as pathogenic. Two of the 22 pathogenic alleles are nonsense mutations that cause autosomal recessive disorders: Hemochromatosis type 2A and Marinesco-Sjogren syndrome. The remaining 18 pathogenic alleles have global minor allele frequencies at least 1% (Table 2). Overall, the CHM1 genome does not appear to harbor an excessive number of risk alleles or extremely rare alleles associated with diseases. (Supplemental data: Clinical Allele Analysis). A similar analysis performed on the GRCh37 assembly identified 3,556 disease susceptibility variants and 15 risk alleles with MAF less than 1% (Chen and Butte, 2011).

### Representation of Segmental Duplication

Analysis of segmental duplication suggests CHM1_1.1 has good representation of large, duplicated sequences. By whole genome assembly comparison (WGAC), we discovered 54,580 pairwise alignments corresponding to 130.9 Mbp of non-redundant duplications or 4.6% of the genome (TableS4). Intrachromosomal events comprise a majority of the segmental duplications with 99.7 Mbp in contrast with 57.3 Mbp of interchromosomal duplications. Additionally, intrachromosomal alignments are generally longer and more similar than interchromosomal alignments (Figure S1). Both of these patterns are consistent with our previous WGAC analysis of GRCh37 (Sudmant et al., 2013). Using an alternative approach to detect segmental duplication based on read depth analysis (WSSD, (Bailey et al., 2001)), we identified 124.6 Mbp of duplicated sequences (4.4% of the genome). These WSSD duplications supported 89.5 of 96.1 Mbp (93%) WGAC duplications that were also >=10 Kbp and >94% identity (Figure S2). Correspondingly, 119.6 Mbp of WSSD duplications (96%) overlapped or occurred within 5 Kbp of a WGAC duplication. To determine how CHM1.1 WGAC duplications compared to duplications from GRCh37, we remapped the WGAC alignments from CHM1.1 to GRCh37.p13 with NCBI’s remap tool. After remapping and omitting coordinates from patches, there were 137.7 Mbp of CHM1.1 duplications. The two assemblies shared 124 Mbp of duplications corresponding to 90% of CHM1.1 duplications and 86% of GRCh37 (Figure 3).

### Identification of misassemblies

The goal for this project is a completely closed reference assembly containing no gaps. Therefore, it is critical for us to identify the extent of misassembly as well as the specific regions involved for targeted correction. We have already begun the process of loading the assembly and curation regions into the GRC curation database and framework. We performed three separate analyses to assess the integrity and identify potential misassemblies.

### Identification of heterozygous SNVs

CHM1 is an essentially homozygous resource. Thus, there should be no heterozygous SNVs identified upon aligning the CHM1 reads to GRCh37, and there should be no SNVs identified when these reads are aligned to the CHM1_1.1 assembly. We were therefore interested in using SNV detection to identify potentially misassembled regions in both GRCh37 as well as CHM1_1.1. First, we aligned the Illumina reads from CHM1 libraries to the GRCh37.p13 primary assembly and identified 99,572 heterozygous sites and 2,445,270 homozygous sites. We stratified heterozygous SNVs based on whether they overlapped repetitive or low complexity sequence (Table S5). A recent study demonstrated that up to 60% of heterozygous SNVs called from CHM1 Illumina reads aligned to the reference are within low complexity regions (LCRs) of the human genome (Li, 2014). We focused on 25,529 heterozygous variants that did not fall within a repetitive sequence (heterozygous non-repetitive: HNR variants), as these may be sites of cryptic duplication in the reference sequence or structural variation in CHM1. The HNR variants were overlapped with the RefSeq annotation (Table S6) and the functional consequences of each variant were predicted (Figure 4).

**Figure 4.**
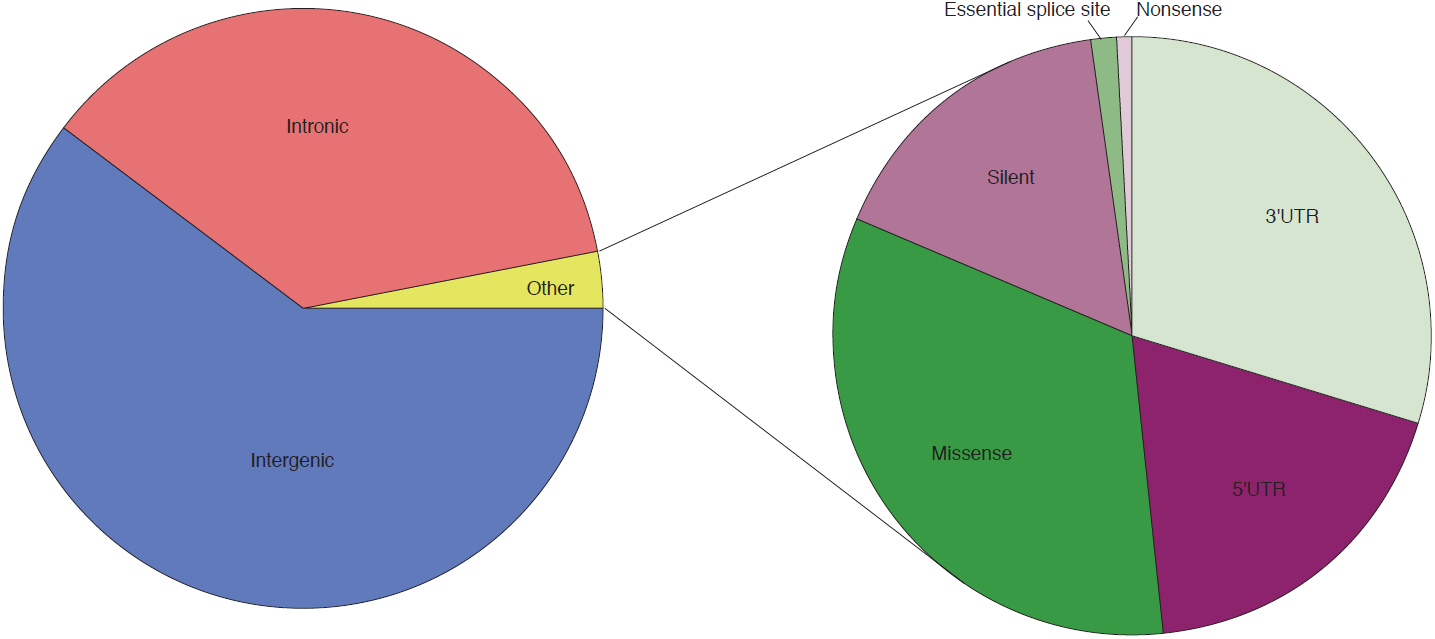
Functional consequences of CHM1 heterozygous variants not in repetitive sequence (HNR variants). Approximately 97% of HNR variants are intergenic or intronic. Of the remaining 3% of other variants, approximately 48% are in the 3′ or 5′ UTR, 17% are silent, and 35% are coding (missense, nonsense, essential splice site).

The genes with the most HNR variants were then compared to genes missing copies in the reference and genes with significantly population stratified copy number (high Vst) from Sudmant et al (Sudmant et al., 2010). Genes with known missing copies in the reference assembly, such as *GPRIN2* and *DUSP22,* have 20 and 56 HNR variants respectively while high Vst genes such as *PDE4DIP* have 267 HNR variants. The gene with the most HNR variants (N=618) is *LOC100996481,* also known as *PRIM2,* that is part of interchromosomal duplications of chromosomes 6 and 3 and represents cryptic segmental duplications in the GRCh37 reference genome (Genovese et al., 2013). Additionally, two regions that were incorrectly represented in GRCh37 and subsequently resolved in GRCh38 using the CHM1 derived BAC library, *SRGAP2* (Dennis et al., 2012) and *IGH* (Watson et al., 2013), both had high counts of HNR variants (39 and 54, respectively) providing additional support for the hypothesis that heterozygous calls are indicative of reference assembly errors. The majority of the heterozygous calls are errors that arise during variant detection due to paralogous sequences mapping to LCRs.

We then aligned the Illumina reads from the CHM1 libraries to the CHM1_1.1 assembly. A total of 86,544 SNVs were called, and 79% of these variants overlap repetitive sequence (RepeatMasker and WGAC; Table S7). There is a significant enrichment of variants in repetitive sequence compared to sequence not annotated as repetitive (1,000 permutations, simulation based p-value < 0.001). Thirty-four regions totaling 49MB have SNV density per kb two standard deviations higher than the mean SNV density per kb of 0.03 (Table S8). Sixty-four percent of the bases in SNV rich regions are annotated as repetitive. There are 294 unique RefSeq and 198 unique Gnomon genes in SNV rich regions including the beta-defensin gene cluster on chromosome 8 and *NBPF1* on chromosome 1 (Table S9). These regions are highly duplicated and the variant calls could represent paralogous sequence variants.

### CH17 BAC ends mapped to CHM1_1.1 assembly

We aligned a set of BAC end sequences derived from the CHORI-17 BAC library to the CHM1_1.1 assembly. As this is the same DNA source as the assembly, there should be no structural variation. The majority of placements were concordant (96.22%), suggestive of a high quality assembly; however, regions with multiple discordant alignments may represent assembly errors. A query set of 306,838 BAC end sequences representing 158,396 unique clones from the CH17 BAC library was aligned to the CHM1_1.1 assembly (Table S10). We identified 1,192 regions with 3,927 unique clones that likely contained assembly errors based on an unexpected size distribution of the aligned BACs. Among unique discordant clones, 2,840 suggested a deletion in the assembly and 443 suggested additional sequence in the CHM1_1.1 assembly not represented in the BAC resource. The regions demonstrating insertion may be due to instability in BAC clones. On average, there are significantly more bases in segmental duplications (WGAC) in the single discordant and multiple mapped clone ends compared to the single concordant clone ends (means=0.24, 0.96, 0.04 and standard deviations=0.18, 0.14, 0.02 respectively for single discordant, multiple and single concordant; Student’s T test, two tailed p=0 for each comparison). The remaining unique discordant placements were comprised of incorrectly oriented ends, indicating that the assembly and clone sequences are inverted relative to one another.

CH17 clones with discordant placements on the CHM1_1.1 assembly may be used to identify regions misassembled due to errors in the reference or genomic variation. For example, in the SMA duplication region at 5q13.3 (NC_018916.2; Figure 5, Figure S3), the GRCh37 reference chromosome represents a single resolved SMA haplotype (Schmutz et al., 2004). However, many CH17 clones aligning to the corresponding region of the CHM1_1.1 assembly have discordant placements that are characterized by inversions and size discrepancies, suggesting that the CHM1_1.1 assembly does not faithfully represent the CHM1 genome at this locus. This observation is consistent with the known variability of this genomic region in the human population, which is associated with its complex segmental duplication structure (Ogino et al., 2004). It should be noted, however, that the clone placements located within the local BAC assemblies in this region are largely concordant, whereas those associated with WGS contigs are discordant. This result demonstrates how the use of sequence from large insert clones can resolve regions too complex for even the reference-guided assembly of WGS contigs. Assembly with additional BAC clones will likely be required to close the existing gaps and fully resolve the CHM1 genomic sequence in these complex regions.

**Figure 5.**
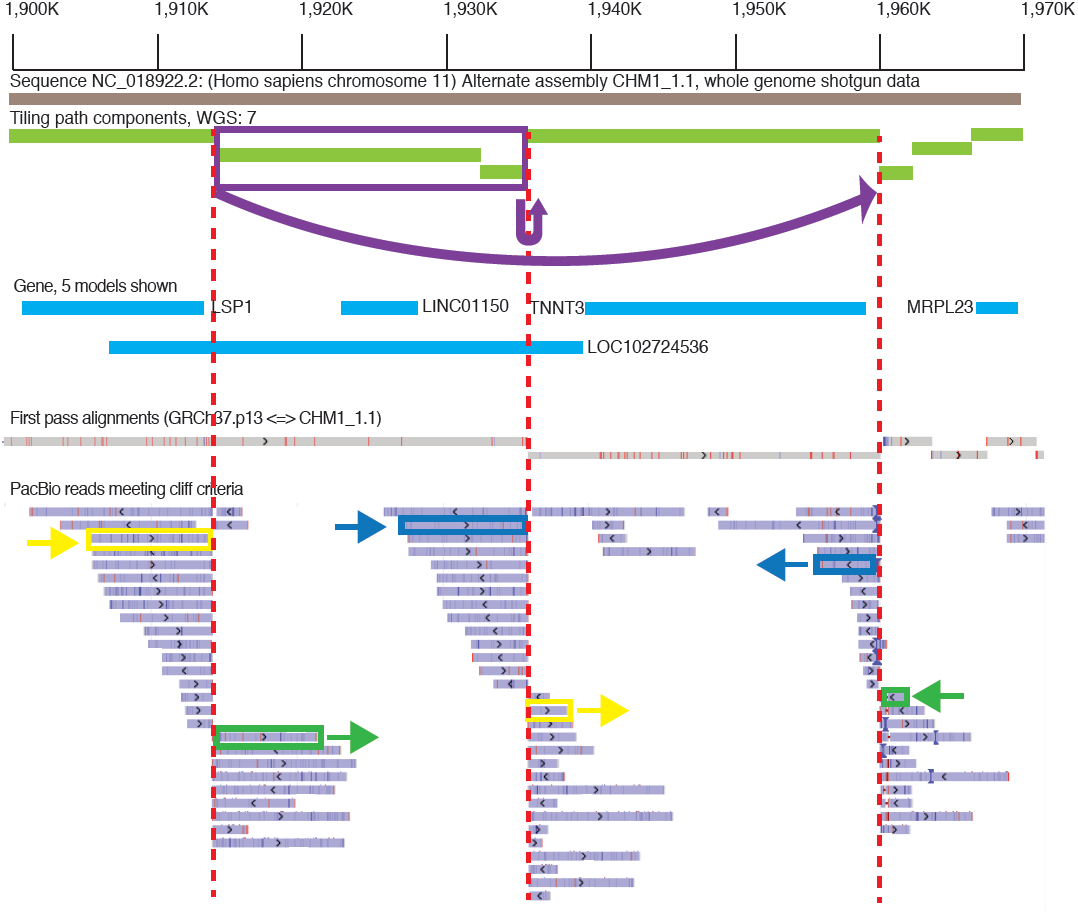
Overview of the NC_018922.2 1.9Mb region, exhibiting three alignment bins with a large number of "cliff" reads. WGS component boundaries flanked by such reads are marked with red dashed lines. Pairs of alignments corresponding to 3 different PacBio reads are marked in yellow, green and blue. These alignments overlap by <10% on each the reads. The split alignments for these 3 reads suggest that the two WGS components marked in purple should be inverted and translocated as indicated by the arrow at the top of the image. The other PacBio reads in these bins exhibit the same pattern of split alignments, which supports the proposed reordering and orientation of the WGS components.

### Alignment to a long read data set for CHM1

To identify errors in the CHM1_1.1 genome assembly (GCF_000306695.2) introduced as a consequence of errors in the GRCh37 primary assembly unit that was used to guide its assembly, we aligned CHM1 PacBio reads (http://datasets.pacb.com/2014/Human54x/fast.html) to the CHM1_1.1 assembly. We hypothesized that these alignments in such regions of CHM1_1.1 would exhibit one or more of the following characteristics: 1) low coverage with respect to coverage in surrounding regions, 2) sharp boundaries at which alignment coverage drops off, or 3) inversions. Low coverage is often associated with highly fragmented assembly regions, which are themselves hallmarks of assembly problems (though they may not necessarily reflect errors introduced by GRCh37). Sharp boundaries could occur at component boundaries (indicative of GRCh37 tiling path errors) or within assembly components (indicative of component assembly errors in GRCh37). Although other assembly features (i.e. repeats or structural variation) can also result in read alignments having similar characteristics, such regions should be enriched for assembly errors.

To identify CHM1_1.1 assembly errors corresponding to unrecognized GRCh37 errors, we focused on CHM1_1.1 assembly sites where alignment coverage dropped off sharply. To this end, we produced a list of regions where there were PacBio aligned reads that met the above criteria, and we refer to these reads as "cliffs". We focused on bins where the cliff count is greater than or equal to ten and the depth is less than 2x the coverage (<108) to eliminate artifacts from repetitive elements. There are 274 loci where cliffs are within 1 kbp of the component boundary and 2109 loci where cliffs are > 1 kbp from component boundary (Supplemental Data: PacBio). Using this approach we are able to clearly visualize regions with assembly errors such as the one on chromosome 11, where two tiling path components are inverted in the CHM1_1.1 assembly and require correction (Figure 5).

## DISCUSSION

There has been a dramatic decrease in sequence cost with a concomitant increase in throughput leading to the availability of thousands of sequenced genomes and exomes. However, analysis of individual genomes depends upon the availability of a high quality reference assembly. Despite the high quality of the human reference assembly, many groups have described shortcomings of this resource, including remaining gaps, single nucleotide errors or gross misassembly due to complex haplotypic variation (Chen and Butte, 2011; Doggett et al., 2006; Eichler et al., 2004; Genomes Project et al., 2012; Kidd et al., 2010). Both gaps and misassembled regions often arise because the DNA sequence used for the assembly was from multiple diploid sources containing complex structural variation. Because such loci often contain medically relevant gene families, it is important to resolve variation at these sites, as the structural and single nucleotide diversity is likely associated with clinical phenotypes (Eichler et al., 2004). Thus, to resolve structurally complex regions and provide a more effective reference resource for such loci, we combined WGS data and BAC sequences from a haploid DNA source to create a single haplotype assembly of the human genome.

Haplotype information is critical to interpreting clinical and personal genomic information as well as genetic diversity and ancestry data, and most previously sequenced individual human genomes are not haploresolved. The current reference human genome sequence represents a mosaic that further complicates haplotyping; within a BAC clone there is a single haplotype representation, but haplotypes can switch at BAC clone junctions. By utilizing an essentially haploid DNA source, we resolved a single haplotype across complex regions of the genome where the reference genome contained a mixture of haplotypes from various sources and/or contained unresolved gaps. For example, a gap on chromosome 4p14 in GRCh37 (chr4:40296397-40297096) was completely resolved using CHM1 WGS data. The gap was flanked by repetitive elements that were not traversed by a clone. This region has subsequently been updated with a complete tiling path in GRCh38.

The addition of high quality BAC sequence to our assembly was vital to resolving gaps. For example, in GRCh37 at chromosome 15q25.2 there was a 79 kbp gap due to over-expansion of a hypervariable region. This region contains many *GOLGA6L* core duplicon genes (Jiang et al., 2007) and highly identical segmental duplications. RP11 BAC clones on one side and RP13 BAC clones on the other side flanked the gap. Using the BAC-based sequence resolved CH17 haplotype, and the gap was filled in GRCh38 (Figure S4). A preliminary analysis of PacBio data shows this region remains unresolved even using long read sequencing. This underscores the importance of curation and employing multiple sequencing strategies to obtain an accurate genome representation.

Despite the high quality of the CHM1_1.1 assembly, we did identify regions that require further improvement. Some of these problems are due to the repetitive nature of the loci, while others are due to using GRCh37 to guide the CHM1_1.1 assembly. The availability of diverse, assembly independent resources, including the recently released long read data set from PacBio provide a pathway for problem identification and correction. The GRC has established the infrastructure to support assembly curation and the development of highly refined reference assemblies, as evidenced by the release of two successive human genome assemblies (GRCh37 and GRCh38) and a mouse genome assembly (GRCm38). We have already begun using these resources to improve the CHM1_1.1 assembly.

We chose a reference-guided assembly method rather than performing a *de novo* assembly of the short WGS reads. An analysis of a *de novo* assembly from short reads using the SOAP algorithm found significant contamination and missing sequences (Alkan et al., 2011). In general, the *de novo* assemblies were approximately 16% shorter than the reference genome, and over 99% of previously validated duplications were missing translating to over 2300 missing coding exons. Another human assembly from massively parallel sequences using the ALLPATHS-LG showed improvements over the SOAP assembly but still only covered ∼40% of segmental duplications. As described above, the gene and repeat coverage of the CHM1_1.1 assembly is comparable to GRCh37. We did not do a formal comparison to GRCh38 because many of the CHM1 BAC tiling paths are used in both assemblies, meaning they are no longer completely independent. Approximately 29Mb of clone sequence and 134kbp of WGS sequence from the CHM1_1.1 assembly has been incorporated into the GRCh38 primary assembly while over 13Mb of clone sequence has been utilized for alternative sequence representations. The somewhat fragmented nature of the CHM1_1.1 assembly means it is not ready to become the Primary assembly in the GRCh series of reference assemblies; however our goal is to improve this assembly so that it could serve this role.

A single haplotype reference assembly will not be sufficient for alignment and variant detection in large-scale human genomic studies. Two individual-specific sequences between a random pair of human individuals ranges between 1.8 and 4Mb (Li et al., 2010). The GRC formalized the concept of multi-allelic representation of complex genome regions with the release of GRCh37. The newest reference genome GRCh38 contains 261 alternative sequence representations at 178 regions, many of which were resolved using the CHM1 data. A recent paper provides the basis for a more formal graph representation (http://arxiv.org/abs/1404.5010) but a great deal of tool development needs to occur before we can formally move to such an assembly representation. While this development occurs, the current multi-allelic reference provides data that allow us to explore complex genomic regions. The use of the single haplotype CHM1 resource has proven quite valuable in resolving several complex regions of the human genome. In many of these cases, the GRCh37 representation was the mixture of several haplotypes and not likely found in any individual. We plan on continuing to develop this resource in an effort to ensure that we have at least one correct representation of all loci in the human genome.

## METHODS

### Cell Line

CHM1 cells were grown in culture from one such conception at Magee-Womens Hospital (Pittsburgh, PA) after parental consent and IRB approval. Cryogenically frozen cells from this culture were grown and transformed using human telomerase reverse transcriptase (hTERT) to develop a cell line. This cell line retains a 46,XX karyotype and complete homozygosity. It was subsequently used for genomic research by multiple investigators and was also used to prepare a BAC library (CHORI17; https://bacpac.chori.org/) for further research.

### Illumina Sequencing

We performed whole genome shotgun sequencing on the CHM1 DNA. KAPA qPCR was used to quantify the libraries and determine the appropriate concentration to produce optimal recommended cluster density on a HiSeq2000 V2 or V3 2x100bp sequencing run. HiSeq2000 V2 and V3 runs were completed according the manufacturer’s recommendations (Illumina Inc, San Diego, CA). We generated over 617 Gb of sequence used for the assembly. The average insert size was 315 bp for 3 libraries, 3 kbp for 3 libraries and 8 kbp for 2 libraries.

### Assembly

Assembly of CHM1 genome used deep coverage WGS sequence reads generated using the Illumina HiSeq platform. This data is publicly available in NCBI’s sequence read archive under project SRP017546. The project has nine experiments of which one was a pilot experiment using 25 bp unpaired reads while remaining eight were all paired-end reads. These eight experiments had a total of 31 runs and were used in producing the assembly (Table Sll).

Reads were aligned to GRCh37 primary assembly using SRPRISM v2.3 aligner. A reference-guided assembly was produced using ARGO vl.O. This assembly has 2,818,728,129 bp in 47737 contigs with N50 of 139647 bp. Both SRPRISM and ARGO were developed at NCBI, but are not yet published. SRPRISM v2.5 is available at ftp://ftp.ncbi.nlm.nih.gov/pub/agarwala/srprism/. Briefly, SRPRISM creates an index on the reference genome and uses the index to find locations on the genome to do extensions. It has resource requirements and performance characteristics comparable to the fastest available aligners, yet provides explicit criteria for search sensitivity and reports all results that have the same quality. ARGO uses conservative heuristics that take into account the insert size and read orientation to produce a most likely sequence for the assembly.

The second source of information provided for the assembly was clones that were specifically designed to address repeat regions. Two hundred ninety six clones in 45 tiling paths and 104 singleton clones were provided. By mapping the clone information to the reference-guided assembly using BLAST and manually reviewing the alignments to decide the best location in the assembly to incorporate the clone sequence, some of the worst regions of the assembly were significantly improved. Four singleton clones could not be used as they are significantly diverged from the assembly. Fourteen additional clones were redundant with other clones. Clone AC243629.2 has an internal expansion that was discovered after the assembly release and has now been subsequently removed. After incorporating clone information, the assembly had 2,846,046,639 bp in 41,406 contigs with an N50 of 143,718 bp. Prior to submission of the assembly to GenBank, the contigs were subsequently filtered to remove some WGS that was redundant to one of the clone paths, to remove small WGS contigs at chromosome termini, to trim terminal Ns from WGS contigs and to accommodate a newly finished clone component and then scaffolded according to alignment with the GRCh37 primary assembly.

### Gene Annotation

The CHM1_1.1 assembly was masked using RepeatMasker and annotated using the NCBI Eukaryotic Genome Annotation Pipeline. Briefly, the assembly is masked using RepeatMasker and then aligned to a set of same-species RefSeq transcripts and genomic sequences to directly annotate the gene, RNA and CDS features. The assembly is also aligned to Gnomon gene prediction models. Gnomon is a two-step gene prediction program that assembles overlapping alignments into "chains" followed by a prediction step that extends the chains into complete models and creates full *ab initio* models, using a Hidden Markov Model (HMM). If the RefSeq and Gnomon models are predicted to have the same splice pattern, the RefSeq transcripts are given precedence. Gnomon predictions are included in the final set of annotations if they do not share all splice sites with a RefSeq transcript and if they meet certain quality thresholds.

### Segmental Duplication Annotation

We applied whole-genome assembly comparison (WGAC) and read depth CNV (WSSD) methods to discover segmental duplications in the CHM1.1 reference assembly. For WGAC analysis, we eliminated all repetitive sequences from the assembly as annotated by RepeatMasker, identified alignments greater than 1 Kbp and with higher than 90% identity, and refined alignments into pairwise duplication calls as previously described (Bailey et al., 2001). Duplication and RepeatMasker files are in Supplemental Data: Duplication Analysis

For read depth CNV analysis, we aligned Illumina whole-genome shotgun (WGS) reads from 11 lanes (SRR642629, SRR642634, SRR642635, SRR642638, SRR642639, SRR642640, SRR642641, SRR642642, SRR642643, SRR642683, SRR642746) with mrsFAST (v. 2.5.0.4) (Hach et al., 2010) and called raw copy number across 1 Kbp windows as previously described (Sudmant et al., 2010). From these raw copy number calls, we identified duplications as regions with copy number >= 3 and >=10 Kbp of non-repeat, non-gap sequence.

### Assembly-Assembly Alignment

We aligned the CHM1_1.1 assembly to the GRCh37 and HuRef assemblies using the two-phase NCBI pipeline. Aligning the two assemblies using BLAST generates the first phase alignments and any locus on the query assembly must have 0 or 1 alignment to the target assembly. Additionally, we use in-database masking through precomputed WindowMasker masked regions. BLAST alignments are then trimmed and post-processed to remove low quality and spurious alignments. Chromosome to chromosome alignments are favored over chromosome to scaffold or scaffold to scaffold alignments. Alignments based on common components are then merged into the longest, consistent stretches possible resulting in a set of alignments called the ‘Common component set’. We then eliminate remaining BLAST alignments that are redundant with the common component alignments. The remaining alignments are then merged independently of the common component alignments and redundant alignments are removed. The two alignment sets are then combined into a single set of alignments and then sorted to select the ‘First pass set’, which are ranked to favor, in order: 1) common component alignments, 2) chromosome to chromosome alignments, 3) alternate to alternate alignments, 4) chromosome to alternate alignments, and 5) count of identities. Finally, only alignments with non-conflicting query/subject ranges are kept for the First Pass set. Conflicting alignments are reserved for evaluation in the ‘Second Pass’. In order to capture duplicated sequences, we do a ‘Second Pass’ to capture large regions (>1Kb) within an assembly that have no alignment, or a conflicting alignment, in the First Pass. In the ‘Second Pass’ alignments, a given region in the query assembly can align to more than one region in the target assembly.

### Variant Analysis

For variant analyses, Illumina reads from CHM1 genomic DNA were mapped to the GRCh37 primary assembly reference using BWA version 0.5.9. Single nucleotide variants (SNVs) were called using both SAMtools and VarScan v2.2.9. Variants were filtered to remove false positives due to alignment and sequencing errors using the values in Table S12. The Illumina reads were then aligned to the CHM1_1.1 assembly and variants were called using the same parameters as above. We overlapped the variants with RefSeq and Gnomon gene annotations as well as segmental duplications (WGAC) and RepeatMasker. SNV density per kilobase and transition:transversion ratio (Ts:Tv) were calculated in 1MB non-overlapping windows using vcftools version 0.1.11.

### BAC end sequence mapping

BAC end sequences from the CH17 BAC library generated from the CHM1 cell line were aligned to the CHM1_1.1, GRCh37 and GRCh38 assemblies and clone placements generated as described in (Schneider et al., 2013). BAC end mappings are provided in Supplemental Data: BAC end mapping. On the CHM1_1.1 assembly, the average insert length = 208,638 and the standard deviation = 20,197. On GRCh37, the average insert length = 208,637 and the standard deviation = 20,149. BAC ends from single concordant, single discordant and multiply mapped clones were evaluated for segmental duplication content and overlapped with gene annotation from RefSeq and Gnomon from the CHM1_1.1 assembly.

### Clinical allele analysis

We obtained data from the NHGRI GWAS catalog using the UCSC browser track intersected with the dbSNP137 track. If the risk allele was reported on the negative strand, we were able to use the dbSNP137 information to correctly assign risk alleles to the positive strand. Additionally, we downloaded the vcf file containing the ClinVar data from NCBI. We took the unique union of risk alleles from both sources and remapped them to CHM1_1.1 coordinates using NCBI default remap parameters. If there were two or more locations we chose the preferred mapping or discarded both. We then compared the risk allele at each locus with the CHM1 genotype.

### PacBio Alignment

We obtained CHM1 reads from the Pacific Biosciences website (http://datasets.pacb.com/2014/Human54x/fast.html) and aligned them to the CHM1_1.1 assembly using BLASR with the following parameters (-nproc 4-sam-clipping soft-bestn 2-minMatch 12-affineAlign-sortRefinedAlignments). To call cliff regions we required that a PacBio read must have two and only two alignments on CHM1_1.1, both alignments must be on the same CHM1_1.1 sequence, and one of the two alignments must meet the criteria of "Score <-2.0*ReadLength". We also required query coverage of the smaller of two segments be ≥ 10% and that the smaller alignment must still involve at least 10% of the PacBio read. Two alignments could not overlap each other by more than 10% and unique coverage ≥ 50%. Coverage drop-offs that occurred within 1 kbp of a CHM1_1.1 boundary were flagged.

The PacBio reads used for this analysis aligned to the CHM1_1.1 assembly at an average coverage depth of 54x. As expected, coverage at regions containing repetitive sequence was notably higher. To improve our likelihood of detecting examples of mis-assemblies, we restricted our review of this list to sites where surrounding coverage did not indicate the presence of repetitive sequence and the drop-off in coverage was roughly equivalent to surrounding coverage.

## DATA ACCESS

All sequence, assembly and clone data is available through Genbank. Supplementary materials and datasets are available on figshare account: http://dx.doi.org/10.6084/m9.figshare.1091429

## ACKNOWLEDGEMENTS

We thank Pieter de Jong for the creation of the CHORI-17 BAC library used extensively in this project. We would like to acknowledge Nathan Bouk for his expertise in sequence alignment and insightful discussions of alignment data. We would also like to acknowledge the efforts of the production and finishing groups at The Genome Institute, particularly Aye Wollom, Susie Rock, Milinn Kremitzki and Derek Albrecht. E.E.E. is an investigator of the Howard Hughes Medical Institute. This work was supported by NIH Grants 2R01HG002385 & 5P01HG004120 to E.E.E.

